# Comment on “Inverted repeats in the monkeypox virus genome are hot spots for mutation”

**DOI:** 10.1101/2025.08.19.670320

**Authors:** Alexandra V. Madorskaya, Fedor M. Kazanov, Gennady V. Ponomarev, Anna Smolnikova, Daria Garshina, Evgenii V. Matveev, Ruslan Kh. Abasov, Dmitry N. Ivankov, Mikhail S. Gelfand, Marat D. Kazanov

## Abstract

Recent work proposed that APOBEC-signature mutations in monkeypox virus (MPXV) are enriched in cruciform structures formed by inverted repeats. We independently reanalyzed the same dataset with the published settings and several complementary approaches and found no statistically supported enrichment of mutations in these structures. While a modest trend at TC motif at 3’ end of hairpin loop contexts cannot be excluded, current evidence is insufficient. Additional data and careful handling of overlapping structures are needed to clarify any association.

## Introduction

Recently, Dobrovolna *et al*.^1^ reported that APOBEC-signature mutations occurred during the 2022 monkeypox virus (MPXV) outbreak are enriched in predicted cruciform structures of the viral genome formed by inverted repeats (IRs) generating two opposite hairpins. While the preference of APOBEC mutations for specific secondary structures formed in double-stranded DNA has been reported before ^2^, we believe that the statistical support for this conclusion in the case of MPXV is insufficient and merit further examination.

Indeed, the authors based this conclusion on the observation that *“63*.*9% of the SNPs were found to arise within IRs”* ^1^. However, when replicating the cruciform predictions by detecting inverted repeats using the same bioinformatic tool and parameter set described in the study, we found that IRs span 52.4% of the MPXV genome. This implies that even random mutations would be expected to overlap predicted cruciform structures approximately half the time. Moreover, the authors’ calculation of 63.9% (39 out of 61 mutations) included seven mutations located merely ‘close’ to, but not within, the IRs, which may substantially bias the statistical interpretation. When applying the binomial test to the enrichment reported in the original study, specifically, 32 out of 61 mutations occurring within IRs, the resulting p-value is 0.55, indicating no statistically significant evidence to reject the null hypothesis of a random distribution of mutations across the MPXV genome.

We would also like to highlight additional concerns regarding the authors’ approach, which complicates interpretation of the findings. The analysis treats entire cruciform as uniform structures, whereas previous studies have shown that APOBEC-induced mutations are predominantly enriched in hairpin loops, particularly at cytosines located at the 3′ terminus, both within and outside the TC motif ^3^. Moreover, many detected IRs overlap and represent mutually exclusive cruciform structures, making their simultaneous presence *in vivo* unlikely and further limiting biological interpretability.

## Results

To address these issues and rigorously evaluate the authors’ hypothesis, we performed an independent analysis using the same set of MPXV mutations and the same PalindromAnalyzer predictions described in the original paper. In addition to analyzing the unfiltered set of overlapping IRs, we implemented four alternative strategies to resolve IR overlaps and consider biologically plausible subsets of cruciform structures: (1) retaining only one most stable cruciform within each overlapping IRs group; (2) retaining non overlapping cruciforms with the highest stability scores; (3) retaining non overlapping cruciforms that collectively maximize total length (representing the upper bound of genomic coverage); (4) retaining the single shortest cruciform per overlapping group (representing the lower bound of coverage). These approaches allowed us to examine both biologically plausible configurations (strategies 1 and 2) and extreme scenarios of genome coverage (strategies 3 and 4).

We quantified mutation enrichment using two complementary approaches: (i) Monte Carlo simulations, in which mutation positions were randomized across the MPXV genome (preserving the mutation count) and enrichment was recalculated over 100,000 iterations to generate empirical p values; and (ii) binomial testing, comparing the observed number of mutations within predicted cruciforms or hairpin loops to the expected number based on the genome coverage. Analyses were performed separately for (a) entire cruciform structures, (b) hairpin loops, (c) cytosines within TC motifs located at the 3′ end of loops, the canonical APOBEC hotspot, and (d) cytosines at the 3’ end of loops irrespective of the TC motif (extended APOBEC signature^3^).

Our results are summarized in Figure 1. Across all four cruciform selection strategies, as well as the original, unfiltered IRs set, we observed no statistically significant enrichment of mutations within either entire cruciform or hairpin loops. Both simulation-based analyses and binomial testing yielded p-values above 0.01. After correction for multiple testing using the Benjamini–Hochberg procedure, the adjusted p-values were correspondingly higher and exceeded the conventional 0.05 significance threshold, with results remaining non-significant.

**Figure 1.**
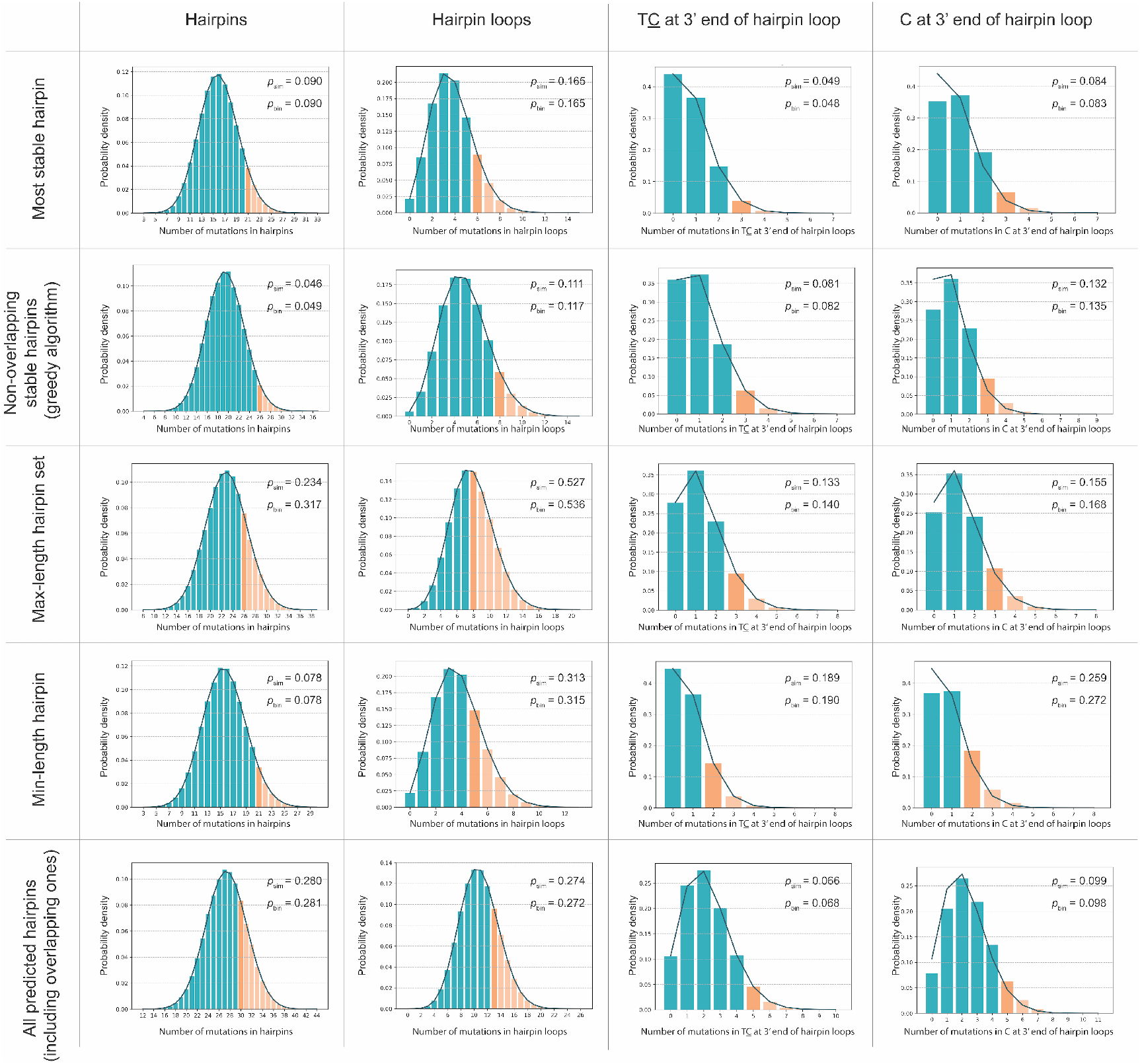
Results of statistical analysis assessing the enrichment of mutations within cruciform structures and APOBEC hotspots located at the 3′ end of hairpin loops. Enrichment was estimated using Monte Carlo simulations (psim) and binomial distribution approximation (pbin), applied to the original set of predicted cruciforms formed by inverted repeats (IRs) and to four sets of non-overlapping IRs selected using different strategies. For each IR set, the mutation enrichment was assessed in the full hairpin structure, the hairpin loop, and TC/C motifs at the 3′ end of the loop (known APOBEC hotspots). The actual number of mutations observed within each considered target region and the corresponding tail of the null distribution are indicated in red.

## Discussion

Notably, a slight trend toward enrichment was observed for mutations in TC motifs at the 3′ end of hairpin loops; however, the number of mutations in this subset is too small to draw firm conclusions. Additional mutation data will be required to evaluate this trend with adequate statistical power. In conclusion, while the hypothesis of APOBEC enrichment in cruciform structures is biologically plausible and supported by analogous findings in human cancers ^4^, our re analysis indicates that the currently available MPXV mutation dataset does not provide statistically significant evidence for this enrichment.

## Methods

The MPXV reference genome (accession GCF_014621545) was downloaded from GenBank. Inverted repeats (IRs) were detected with PalindromAnalyzer ^5^ using the parameters of the original study: IR arm length 0–30 bp, spacer length 0–10 bp, and ≤1 mismatch per arm. Each IR was represented by its outer genomic span (from the earliest arm start to the latest arm end). We then constructed an interval-overlap graph in which nodes are IRs and an edge connects two IRs whose spans overlap by ≥1 bp. Overlap groups were defined as the connected components of this graph.

Within each overlap group, we applied four strategies: (1) select the single cruciform with the lowest predicted free energy (as reported by PalindromAnalyzer output ^6^) as the most-stable representative; (2) apply a greedy non-overlapping procedure that iteratively selects the most stable cruciform and removes all overlapping candidates until none remain; (3) use dynamic programming to choose cruciforms (or hairpin loops, depending on the experiment) that maximize total genomic coverage within the group; and (4) select a single representative that minimizes genomic coverage by the cruciform itself (or by its loop, as appropriate). For binomial approximations, the probability of success parameter *0* was defined as the fraction of the genome covered by the selected cruciforms (or loops), i.e., total covered ba<u>s</u>e pairs divided by the genome length.

## Author contributions

A.V.M., F.M.K., and G.V.P. performed the simulations and statistical analyses. A.S., D.G., E.V.M., and R.K.A. contributed to data preparation and analysis. D.N.I., M.S.G., and M.D.K. contributed to data analysis, wrote and revised the manuscript.

## Acknowledgements

This study was supported in part by RSF grant no. 22-14-00132 (to D.N.I.), which funded the Monte Carlo simulations, and by assignment FFRW-2024-0004 (to G.V.P., E.V.M., and R.K.A.). We thank Irina Ponomareva for designing the preprint layout.

## Data availability statement

All scripts used in this study are publicly available in the GitHub repository: https://github.com/KazanovLab/APOBEC3-Mpox-IRs

